# Rust fungal effectors mimic host transit peptides to translocate into chloroplasts

**DOI:** 10.1101/019521

**Authors:** Benjamin Petre, Cécile Lorrain, Diane G.O. Saunders, Sébastien Duplessis, Sophien Kamoun

## Abstract

Parasite effector proteins target various host cell compartments to alter host processes and promote infection. How effectors cross membrane-rich interfaces to reach these compartments is a major question in effector biology. Growing evidence suggests that effectors use molecular mimicry to subvert host cell machinery for protein sorting. We recently identified CTP1 (<u>c</u>hloroplast-<u>t</u>argeted <u>p</u>rotein 1), a candidate effector from the poplar leaf rust fungus *Melampsora larici-populina* that carries a predicted transit peptide and accumulates in chloroplasts. Here, we show that the CTP1 transit peptide is necessary and sufficient for accumulation in the stroma of chloroplasts, and is cleaved after translocation. CTP1 is part of a *Melampsora*-specific family of polymorphic secreted proteins whose members translocate and are processed in chloroplasts in a N-terminal signal-dependent manner. Our findings reveal that fungi have evolved effector proteins that mimic plant-specific sorting signals to traffic within plant cells.

## INTRODUCTION

Parasitic microbes deliver effector proteins into host cell compartments to manipulate a variety of processes and promote parasitic colonization (Dodds and Rathjen, 2010; Win *et al.*, 2012). One driving theme in plant pathology is to uncover how effector proteins traffic to these compartments; most notably how they cross a multitude of membrane-bound biological interfaces. Bacterial effectors cross the host plasma membrane through a complex molecular syringe (or secretion system), whereas effectors of filamentous parasites, namely fungi and oomycetes, enter host cells via mechanisms that remain to be discovered (Petre and Kamoun, 2014). Once inside host cells, some effectors subsequently traffic to distinct compartments including nuclei, plasma membrane, endoplasmic reticulum, tonoplast, vesicles, chloroplasts and mitochondria (Lindeberg *et al.*, 2012; Deslandes and Rivas, 2012; Hicks and Galan, 2013). To do so, several effectors possess domains that mimic host-targeting (or addressing) sequences. For example, effectors targeted to host nuclei through nuclear pores carry nuclear-localisation signals (NLS) (Schornack *et al.*, 2010; Wirthmueller *et al.*, 2014). However, how pathogens target effectors to membrane-bound organelles such as chloroplasts is poorly understood.

Chloroplasts are plant cell organelles composed of a double membrane envelope that surrounds a fluid called stroma, in which structures such as thylakoids and starch granules reside. The envelope can form elongated structures called stromules (*strom*a-containing tub*ules*), which are thought to improve exchange of molecules with other compartments (Krenz *et al.*, 2012). Chloroplasts selectively internalize proteins from the cytosol via the TOC/TIC (Translocon at Outer membrane of Chloroplasts/Translocon at Inner membrane of Chloroplasts) complex, by recognizing transit peptides that are present at the N-termini of proteins (Shi and Theg, 2013). Upon translocation, transit peptides are typically cleaved off by processing peptidases and mature proteins are released in the stroma (Texeira and Glaser 2013). Transit peptides are variable in term of amino acid sequence and length, and no consensus has been identified (Demarsy *et al.*, 2014).

To date, only a few plant pathogen effectors have been reported to target chloroplasts (Jelenska *et al.*, 2007; Rogriguez-Herva *et al.*, 2012; Li *et al.*, 2014; Petre *et al.*, 2015). HopI1, HopN1, HopK1 and AvrRps4 are type-III effector proteins of the bacterial plant pathogen *Pseudomonas syringae* pv *tomato* that function as virulence factors by modulating chloroplast structures and functions. How HopI1 and HopN1 translocate into chloroplasts is unknown, as they lack a N-terminal transit peptide. In contrast, HopK1 and AvrRps4 carry N-terminal cleavable regions that are sufficient for import into chloroplasts, suggesting that these proteins carry transit peptides (Li *et al.*, 2014). However, whereas both proteins accumulate in chloroplasts in stable *A. thaliana* transgenics, they diffuse in the cytosol and the nucleus when transiently expressed in *N. benthamiana* leaf cells (Li *et al.*, 2014). Besides, several studies have reported AvrRps4 in the nucleus and cytosol rather than, or in addition to chloroplasts (Heidrich *et al.*, 2011; Bhattacharjee *et al.*, 2011; Sohn *et al.*, 2012).

Rust fungi (Basidiomycetes, Pucciniales) are obligate biotrophic parasites of plants (Duplessis *et al.*, 2015). Several rust species are devastating pathogens of crops, and a constant threat to agrosystems worldwide (Pennisi 2010). For instance, the leaf rust fungus *Melampsora larici-populina* triggers annual epidemics on cultivated poplars and dramatically reduces plantations yield (Gérard *et al.* 2006). We recently took advantage of the predicted effector complement of *M. larici-populina* to perform an expression screen of the most promising candidates in leaf cells of the model plant *Nicotiana benthamiana*. These analyses revealed that the effector MLP107772, renamed chloroplast-targeted protein 1 (CTP1) in the present study, accumulates in chloroplasts (Petre *et al.*, 2015). Interestingly, CTP1 carries a predicted transit peptide of 83 amino acids at its N-terminus, which may explain its ability to translocate into chloroplasts (Petre *et al.*, 2015).

In this study, we further exploited the *N. benthamiana* experimental system to study effector trafficking across the chloroplast envelope. We combined genetic transformation with live cell imaging, cell fractionation and biochemical analyses to demonstrate that CTP1 resides in the stroma of chloroplasts. We then showed that the predicted CTP1 transit peptide is necessary and sufficient for accumulation in chloroplasts, and is cleaved upon translocation. CTP1 is a member of a *Melampsora*-specific effector family. We also provide evidence that other members of the family also enter chloroplasts in a N-terminal-dependent manner. Our data indicate that rust fungi have evolved a class of effector proteins that translocate into chloroplasts by functionally mimicking plant-targeting sequences.

## RESULTS

### CTP1-mCherry accumulates in the stroma of chloroplasts and in mitochondria

We recently reported that the mature form of CTP1 fused to a GFP (CTP1_28-171_-GFP) accumulates inside chloroplasts and mitochondria when it is heterologously expressed in *N. benthamiana* leaf cells (Petre *et al.*, 2015). To test the robustness of this observation, we constructed a fusion protein with a different fluorescent tag (CTP1_28-171_-mCherry) and assayed its localization in *N. benthamiana* leaf cells by confocal microscopy. CTP1_28-171_-mCherry fluorescent signal overlapped with chlorophyll autofluorescence, but also labeled stromules and mitochondria-like bodies (Figure 1). Co-expression of CTP1_28-171_-mCherry and ScCOX4-GFP, a marker of mitochondria (Nelson *et al.*, 2007), showed overlapping signals, confirming the presence of CTP1_28-171_-mCherry in mitochondria (Figure S1). The co-expression of CTP1_28-171_-GFP and CTP1_28-171_-mCherry revealed perfectly overlapping fluorescent signals, indicating that the nature of the fluorescent protein does not affect the localization. Saturation of the fluorescent signals revealed no accumulation in the cytosol and the nucleus, indicating that CTP1_28-171_-fluorescent protein fusions are probably internalized into these organelles (Figure S2).

**Figure 1.**
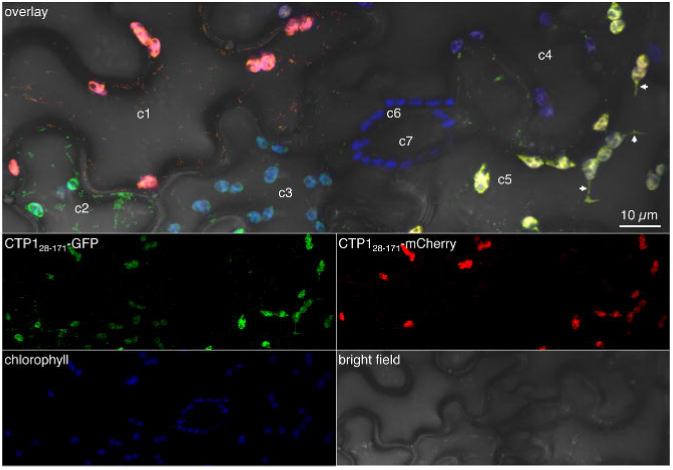
CTP1_28-171_-fluorescent protein fusions traffic to chloroplasts. (A) Live-cell imaging of CTP1_28-171_-GFP and CTP1_28-171_-mCherry in *N. benthamiana* leaf pavement cells. Proteins were transiently expressed in *N. benthamiana* leaf cells by agroinfiltration. Live-cell imaging was performed with a laser-scanning confocal microscope with a sequential scanning mode two days after infiltration. The GFP and the chlorophyll were excited at 488 nm; the mCherry was excited at 561 nm. GFP (green), mCherry (red) and chlorophyll (blue) fluorescent signals were collected at 505-525 nm, 580-620 nm and 680-700 nm, respectively. Images present a maximal projection of 23 optical sections (z-stack: 18.4 μm). White arrows indicate stromules. Cells marked c1 to c5 are pavement cells differentially accumulating the two fusion proteins (c1 and c5 accumulate the mCherry fusion; c1, c2, c3 and c5 accumulate the GFP fusion). Cells marked c6 and c7 are guard cells.

To unambiguously determine whether CTP1 translocates inside chloroplasts, we performed a series of co-localizations of CTP1_28-171_-fluorescent protein fusions with various markers of sub-chloroplastic compartments. First, we co-expressed CTP1_28-171_-GFP with NbpMSRA-mCherry, a marker of the stroma of chloroplasts (Lorrain *et al.*, 2014a). In this assay, GFP and mCherry fluorescent signals overlapped perfectly, indicating that CTP1 was present in the stroma (Figure 2A). Second, co-expression of CTP1_28-171_-GFP with a GBSSI-YFP fusion (a starch granule marker; Wang *et al.*, 2013), allowed a better visualization of chloroplasts at high magnification. The uniform distribution of the GFP fluorescent signal around GBSSI-YFP labeled starch granules further confirmed the accumulation of CTP1_28-_ 171-GFP in the stroma (Figure 2B, Figure S2). Finally, we used cell fractionation to isolate chloroplasts from *N. benthamiana* leaves expressing CTP1_28-171_-GFP. Confocal microscopy performed on purified organelles revealed a strong fluorescent signal in some chloroplasts, which was uniformly distributed around the starch granules (Figure 3). These experiments indicate that CTP1 enters the chloroplasts to accumulate within the stroma.

**Figure 2.**
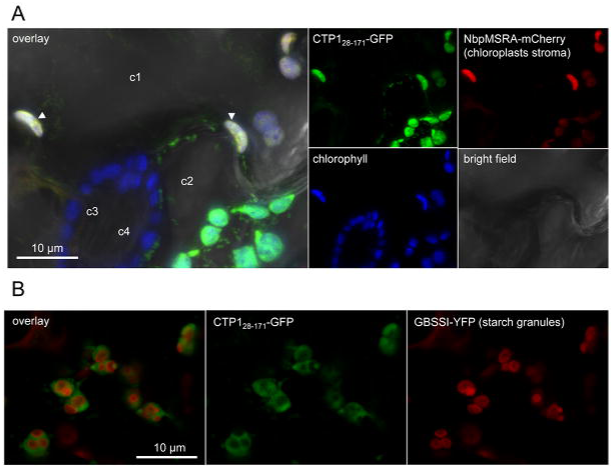
CTP1_28-171_-fluorescent protein fusions translocate in chloroplasts. (A) Live-cell imaging of CTP1_28-171_-GFP and NbpMSRA1-mCherry (chloroplast stroma marker) in *N. benthamiana* leaf pavement cells. Images present a maximal projection of 21 optical sections (Z-stack: 16.8 μm). Cells marked c1 and c2 are pavement cells differentially accumulating the two fusion proteins. Cells marked c3 and c4 are guard cells. White arrowheads show chloroplasts from c1 where GFP, mCherry and chlorophyll fluorescent signals overlap. (B) Live-cell imaging of CTP1_28-171_-GFP and GBSSI-YFP (starch granule marker) in *N. benthamiana* leaf pavement cells. Images present a maximal projection of two optical sections (z-stack: 1.6 μm). Proteins were co-expressed in *N. benthamiana* leaf cells by agroinfiltration. Live-cell imaging was performed with a laser-scanning confocal microscope with a sequential scanning mode two days after infiltration. The GFP and the chlorophyll were excited at 488 nm; the YFP was excited at 514 nm; the mCherry was excited at 561 nm. YFP (red), mCherry (red) and chlorophyll (blue) fluorescent signals were collected at 530-550 nm, 580-620 nm and 680-700 nm, respectively. GFP (green) fluorescent signal was collected at 505-525 nm (A) or 505-510 nm (B).

**Figure 3.**
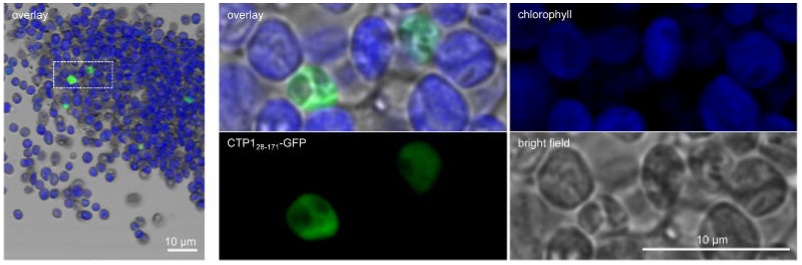
CTP1_28-171_-GFP resides inside isolated chloroplasts. Live-cell imaging of CTP1_28-171_-GFP in chloroplasts isolated from agrotransformed *N. benthamiana* leaf cells. Leaves were homogenized and the lysate was filtered and centrifuged to retrieve organelles. The chloroplasts were separated on a Percoll gradient and collected with a pipet for immediate use. Live-cell imaging was performed with a laser-scanning confocal microscope. The GFP and the chlorophyll were excited at 488 nm. GFP (green) and chlorophyll (blue) fluorescent signals were collected at 505-525 nm and 680-700 nm, respectively. Images present a single optical section. The left-hand panel shows a low magnification overlay image (GFP fluorescence, chlorophyll fluorescence and bright field). The white dotted-line rectangle indicates the close-up presented in the right-hand panels.

### CTP1 carries a functional chloroplast transit peptide

We hypothesized that the predicted 83-amino-acid N-terminal transit peptide of CTP1 mediates trafficking to chloroplasts and mitochondria (Petre *et al.*, 2015; Figure 4A). To test the functionality of this predicted transit peptide, we generated two truncations: CTP1_28-114_ (the predicted transit peptide plus four amino acids), and CTP1_107-171_ (CTP1 without its predicted transit peptide). We fused the truncations with GFP (CTP1_28-114_-GFP and CTP1_107-_ 171-GFP) and determined their subcellular localization in *N. benthamiana* leaf cells. CTP1_28-_ 114-GFP signal overlapped with chlorophyll, but also showed a weak signal in the cytosol and the nucleus (Figure 4B). We used CTP1_28-171_-mCherry as a marker of both the stroma and mitochondria, and co-expressed it with CTP1_28-114_-GFP. Fluorescent signals overlapped in the stroma but not in mitochondria, suggesting that CTP1_28-114_-GFP accumulates only in chloroplasts. Further observations performed on chloroplasts isolated from leaves expressing CTP1_28-114_-GFP confirmed the presence of the fusion protein in the stroma (Figure 4C). On the other hand, CTP1_107-171_-GFP signal was not detected in chloroplasts or mitochondria, and showed a nucleocytosolic distribution (Figure 4D). This indicates that CTP1 lacking its transit peptide can no longer enter these organelles. In stable *N. benthamiana* transgenics, CTP1_28-_ 114-GFP fluorescent signal specifically labeled chloroplasts and stromules, validating the observations made in transient assays (Figure 4E). We conclude that CTP1 carries a functional transit peptide, which is necessary and sufficient to mediate trafficking to chloroplasts. However, this transit peptide is necessary but not sufficient for accumulation in mitochondria.

**Figure 4.**
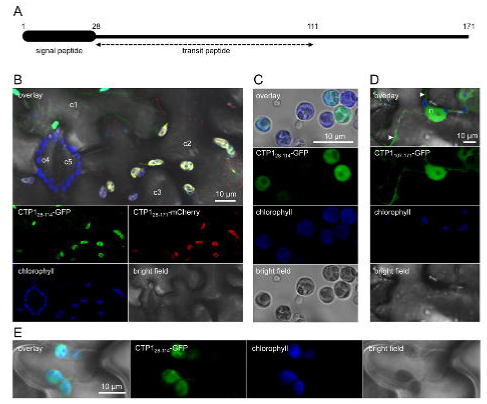
CTP1 carries a functional chloroplast transit peptide. (A) Protein architecture of CTP1, with its two predicted targeting sequences. (B) Live-cell imaging of CTP1_28-114_-GFP and CTP1_28-171_-mCherry (here used as a marker of the stroma of chloroplasts and mitochondria) in *N. benthamiana* leaf pavement cells. Images present a maximal projection of 11 optical sections (z-stack: 8.8 μm). Cells marked c1 to c3 are pavement cells differentially accumulating the two fusion proteins. Cells marked c4 and c5 are guard cells. (C) Live-cell imaging of CTP1_28-114_-GFP in chloroplasts isolated from agrotransformed *N. benthamiana* leaf cells. Leaves were homogenized and the lysate was filtered and centrifuged to retrieve organelles. The chloroplasts were separated on a Percoll gradient and collected with a pipet for immediate use. Images present a single optical section. (D) Live-cell imaging of CTP1_107-171_-GFP in *N. benthamiana* leaf pavement cells. Images present a maximal projection of three optical sections (z-stack: 2.4 μm). White arrowheads indicate cytosol. n: nucleus. Proteins were expressed in *N. benthamiana* leaf cells by agroinfiltration. Live-cell imaging was performed with a laser-scanning confocal microscope with a sequential scanning mode two days after infiltration. The GFP and the chlorophyll were excited at 488 nm. The mCherry was excited at 561 nm. GFP (green) and chlorophyll (blue) fluorescent signals were collected at 505-525 nm and 680-700 nm, respectively. (E) Live-cell imaging of CTP1_28-114_-GFP in leaf pavement cells of a stable transgenic *N. benthamiana* plant. Images present a single optical section.

### CTP1 is part of a *Melampsora*-specific family of modular and polymorphic proteins

*CTP1* was first described as an orphan gene in *M. larici-populina* (Hacquard *et al.*, 2012; Saunders *et al.*, 2012). However, a more recent analysis reported that CTP1 displays sequence similarity to other secreted proteins of *M. larici-populina* (Cantu *et al.*, 2013). To establish a list of proteins related to CTP1, we performed sequence similarity searches as well as protein and gene structure similarity analyses. We identified four proteins with sequence similarity to CTP1: Mlp52246 (hereafter CTP2), Mlp55690 and Mlp116083 in *M*.*larici-populina*, and Melli_sc2834 (hereafter CTP3) in the flax rust fungus *Melampsora lini*. The four proteins showed amino acid similarity to CTP1 that ranged from 16 to 28%. Although the sequence similarity was low, these five proteins display a similar modular protein structure. They all have a predicted signal peptide followed by a predicted transit peptide of variable length, as well as two to three repeated domains or modules, with the first one overlapping with the predicted transit peptide (Figure 5A). Also, gene structures matched protein organization, with the first exon encoding the signal peptide, and each of the other exons encoding a module. These modules are highly divergent (average paired amino acid identity below 30%, ranging from 15 to 100%) but present stretches of amino acids with similar properties and four conserved cysteine residues (Figure 5B). Interestingly, the phylogeny of the modules does not fully match the gene organization (Figure 5B), suggesting that the CTP family evolved potentially by exon shuffling. We conclude that CTP1 is part of a family of secreted, modular proteins that have probably emerged and diversified recently in *Melampsora* species.

**Figure 5.**
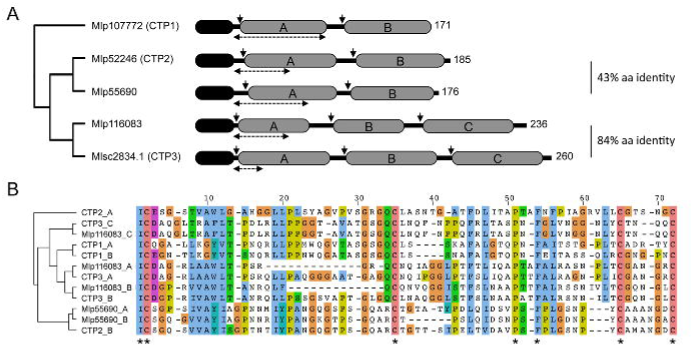
CTP1 is part of a family of modular and polymorphic proteins. (A) Gene and protein architecture of Mlp107772 (alias CTP1), Mlp52246 (alias CTP2), Mlp55690, Mlp116083, and Melli_sc2834.1 (alias CTP3). Black: predicted signal peptide; horizontal double-arrowed dotted lines: predicted transit peptide; vertical arrows: intron positions; grey: repeated domains or modules. Numbers on the right-hand side indicate the number of amino acids for each protein. Chloroplast transit peptides were predicted with the ChloroP 1.1 Server and the following Score/CS-Score/cTP-length values were obtained: CTP1 (0.533/1.040/83); CTP2 (0.503/4.184/38); Mlp55690 (0.484/5.625/67); Mlp116083 (0.510/2.432/40); CTP3 (0.493/1.934/23). (B) Phylogeny and amino acid alignment of the 12 repeated modules displayed in (A). Modules are named according to their protein of origin and their position in the protein sequence (A, B or C) as indicated in (A). The phylogenetic tree was built with Jalview (neighbor joining using % identity) and the amino acid alignment was performed with ClustalX. Amino acid residues are colored according to the ClustalX scheme. Black asterisks mark the seven fully conserved residues.

### CTP2 and CTP3, two proteins similar to CTP1, accumulate in chloroplasts

To determine whether CTP1-like proteins also target chloroplasts and mitochondria, we studied CTP2 and CTP3, two proteins representative of the diversity of the family, and which carry two and three modules, respectively. We generated fusions between the mature forms of CTP2 and CTP3 (i.e. without the signal peptide) with GFP or mCherry and expressed the fusion proteins in *N. benthamiana*. Confocal microscopy revealed that the two fusion proteins display a signal that overlapped with chlorophyll. CTP2 fluorescent signal accumulated in the stroma of the chloroplasts but also in the nucleus and cytosol (Figure 6A, Figure S4). CTP3 fluorescent signal distributed similarly to CTP2, but also labeled discrete areas of approximately 0.1 to 0.3 μm of diameter within the chloroplasts (Figure 6B, Figure S5A and S5B). A *Populus trichocarpa* Coproporphyrinogen III Oxidase-GFP fusion (PtCPO-GFP) was previously observed to accumulate in similar sub-chloroplastic structures in *N. benthamiana* (Lorrain *et al.*, 2014b). Co-expression of CTP3_28-260_-mCherry and PtCPO-GFP revealed overlapping signals, indicating that CTP3_28-260_-fluorescent protein fusions reside in PtCPO-containing compartments whose nature remains to be determined (Figure S5C). Co-expression of CTP2_29-185_- and CTP3_28-260_-fluorescent protein fusions with mitochondria markers (ScCOX4-GFP or ScCOX4-mCherry) revealed no overlapping signals, suggesting that CTP2 and CTP3 do not accumulate in mitochondria (Figure S4, Figure S5A). We conclude that CTP2 and CTP3 translocate into chloroplasts when expressed in *N. benthamiana* cells.

**Figure 6.**
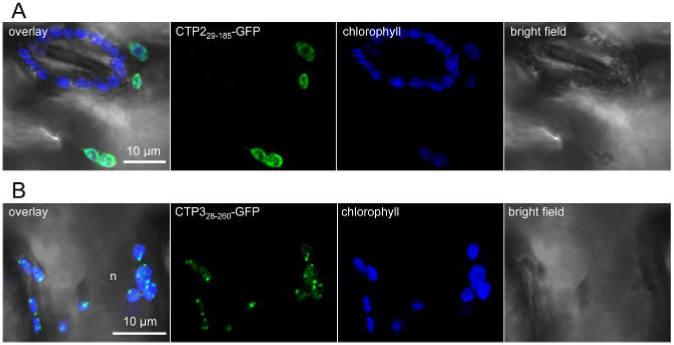
CTP2_29-185_-GFP and CTP3_28-260_-GFP accumulate in chloroplasts. (A) Live-cell imaging of CTP2_29-185_-GFP in *N. benthamiana* leaf pavement cells. Images present a maximal projection of seven optical sections (z-stack: 5.6 μm). (B) Live-cell imaging of CTP3_28-260_-GFP in *N. benthamiana* leaf pavement cells. Images present a maximal projection of 15 optical sections (z-stack: 12.0 μm). Live-cell imaging was performed with a laser-scanning confocal microscope two days after infiltration. The GFP and the chlorophyll were excited at 488 nm. GFP (green) and chlorophyll (blue) fluorescent signals were collected at 505-525 nm and 680-700 nm, respectively.

### CTP1, CTP2 and CTP3 are processed in chloroplasts

Upon translocation in chloroplasts and mitochondria, processing peptidases cleave off transit peptides (Teixeira and Glaser, 2013). To determine whether CTPs undergo processing *in planta*, we performed anti-GFP western blotting with total proteins isolated from *benthamiana* leaves expressing the CTP-GFP fusions. The band signals for all fusion proteins were smaller than their theoretical size, suggesting that they undergo cleavage *in planta*. Indeed, CTP1_28-171_-GFP showed a unique band signal at 34 kDa, approximately 8 kDa below its expected size of 41.8 kDa (Figure 7), confirming previous observations (Petre *et al.*, 2015). Both CTP2_29-185_-GFP and CTP3_28-260_-GFP displayed a band signal that coincided with their predicted molecular weight at 42.7 and 51.0 kDa, respectively, but also a second band signal that was 8 and 16 kDa smaller, respectively. The size of all these band signals coincide with the predicted molecular weight of the last repeat module of each protein fused to GFP, which suggests that cleavage events occur between modules A and B (CTP1 and CTP2) and between modules B and C (CTP3) (Figure 5).

**Figure 7.**
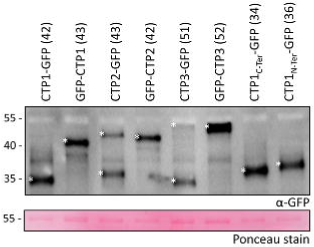
CTP1, CTP2 and CTP3 undergo processing inside chloroplasts. Anti-GFP Western blotting analyses of GFP fusions. Proteins were transiently expressed in *N. benthamiana* leaf cells by agroinfiltration. Total protein extraction was performed two days after infiltration by grinding leaves in liquid nitrogen and immediately reducing and denaturing proteins from the leaf powder in a Laemmli buffer 15 min at 95°C. Proteins were separated by 15% SDS-PAGE and transferred onto a nitrocellulose membrane. Immune detection was performed with anti-GFP antibodies. For each fusion protein the theoretical size is indicated in parentheses in kiloDaltons (kDa). Protein sizes are indicated on the left-hand side of the image in kDa. Asterisks indicate protein bands discussed in the text. Ponceau staining of the RubisCO onto the membrane was used as a loading control.

To determine whether processing events occur specifically in chloroplasts, we generated CTP fusions that are expected to interfere with chloroplast translocation. Since the addition of N-terminal amino acid residues is known to abolish the functionality of transit peptides (Carrie *et al.*, 2009), we constructed N-terminally GFP tagged fusion proteins (GFP-CTP1_28-171_, GFP-CTP2_29-185_ and GFP-CTP3_28-260_) and expressed them in *N. benthamiana* leaves. We did not detect fluorescent signal in chloroplasts with the three fusion proteins, which accumulated only in the nucleus and the cytosol (Figure S6). Each of the three fusion protein yielded a single band signal around its predicted molecular weight with anti-GFP western blot analyses, demonstrating the absence of processing in the nucleus and the cytosol (Figure 7). From this set of experiments, we conclude that CTP1, CTP2 and CTP3 translocate in chloroplasts in a N-terminal signal-dependent manner, and that they undergo processing inside chloroplasts.

## DISCUSSION

The N-terminus of the rust fungal protein CTP1 is necessary and sufficient to enter the chloroplast organelle of plants indicating that it mimics host transit peptides to subvert host cell machinery. Host mimicking domains are frequent in pathogen effectors (Hicks and Galan, 2013). However, how did a chloroplast transit peptide evolve in an organism that lacks chloroplasts? It should be noted that mitochondria and chloroplast transit peptides have similar properties (Carrie and Small 2013). The CTP1 transit peptide may have evolved from a mitochondrial transit peptide endogenous to the fungus. The observation that CTP1 can accumulate in mitochondria somewhat supports this hypothesis although the transit peptide does not fully account for this (Figure 4). On the other hand, the transit peptide may have evolved *de novo*. Transit peptides typically do not display conserved amino acid sequences, and their functionality is based on conserved physical properties (Bruce 2001). Therefore, they are more likely to evolve *de novo* compared with other domains whose function is based on strict amino acid sequence conservation (Tonkin *et al.*, 2008). This possibility is perhaps more likely given that CTP1 does not share sequence similarity to any known protein sequence, and belongs to a polymorphic and lineage-specific family. Hence, the CTP1 transit peptide may have evolved *de novo* possibly by neofunctionalization after a duplication or recombination event.

Host-translocated effectors of oomycete and fungal plant pathogens usually carry host-targeting sequences just downstream of their signal peptide (Petre and Kamoun 2014). These targeting sequences have no known function in host cells. We have shown that the N-terminal region immediately downstream of CTP1 signal peptide functions as transit peptide in plant cells. This finding implies that, if CTP1 encodes a N-terminal host-targeting sequence, it overlaps with its transit peptide. It is also possible that CTP1 does not require a host-targeting sequence.

CTP1 accumulates in both chloroplasts and mitochondria. Although accumulation in mitochondria is relatively weak, CTP1 could very well target the two organelles. Many plant proteins have evolved to target and function in two cell compartments, and are called ‘dually-targeted’ proteins (Carrie and Whelan 2013). For instance, over a hundred plant proteins target both chloroplasts and mitochondria (Carrie and Small 2013). Some pathogen effectors have also been reported in multiple host cell compartments (Heidrich *et al.*, 2011; Hicks and Galan, 2013; Li *et al.*, 2014) and may also be dually-targeted proteins. However, it has to be noted that disparate locations observed by different authors could be artifactual, for instance due to the use of different tags or methods of localization. In rust fungi, several members of the RTP1 family have been immunolocalised in various compartments within the plant cell and at the host-pathogen interface (Kemen *et al.*, 2005; Hacquard *et al.*, 2012; Kemen *et al.*, 2013).

CTP1, CTP2 and CTP3 display various patterns of accumulation in chloroplasts and mitochondria. Plant proteins evolve to form large families, whose members diversify to target and function in different cell compartments (Rouhier *et al.*, 2006). The same holds true for effectors of plant parasites. For instance, several members of the Avrblb2 family of *Phythophtora infestans* accumulate in the nuclei, the cytosol or at the plasma membrane (Bozkurt *et al.*, 2011; Zheng *et al.*, 2014). Also, effector alleles can target different compartments (Leonelli *et al.*, 2011; Caillaud *et al.*, 2012). Whether effector variants carry similar or different functions in distinct cellular compartments remains to be determined. In plants, members of large protein families often carry similar biochemical activities, although they are performed with substrates or protein partners that are specific to a given compartment (Rouhier *et al.*, 2006).

We have shown that three candidate effectors from *Melampsora* species accumulate in chloroplasts (and to some extent in mitochondria) in *N. benthamiana* leaf cells. Growing evidence indicates that effectors from unrelated parasites converge towards chloroplasts and mitochondria (Win *et al.*, 2012; Hicks and Galan 2013; Li *et al.*, 2014; Petre *et al.*, 2015). This implies that manipulating chloroplasts and mitochondria is crucial for parasites to colonize host plants. Surprisingly, to date no oomycete effectors have been reported in these organelles, despite several large-scale subcellular localisation screens (Schornack *et al.*, 2010; Caillaud *et al.*, 2012; Stam *et al.*, 2013; Win *et al.*, unpub). However, all these studies used N-terminally tagged effectors, which may have impaired the proper translocation of the proteins into the organelles, resulting in false negatives.

The ability of CTPs to enter chloroplasts suggests that they have a function inside this organelle. What could be that function? CTPs display no sequence similarity to known proteins or protein domains, which prevents the prediction of a putative function based on their primary structure. The only similar feature among the CTPs is the four fully conserved cysteine residues within repeated modules. Such patterns of conserved cysteine residues have been observed previously for several families of secreted proteins in *M. larici-populina* (Duplessis *et al.*, 2011; Hacquard *et al.*, 2012). Repeated domains with conserved residues are common within effectors (Kloppholz *et al.*, 2011; Mak *et al.*, 2012; Chou *et al.*, 2011) and can provide conserved folds that promote rapid evolution (Win *et al.*, 2012). The cysteines could have different roles, for instance by being involved in intra- or extra-molecular disulfide bridges or by having a catalytic activity. Importantly, cysteine residues are also conserved in the repeats that overlap with transit peptides. It is tempting to speculate that repeats could be independently functional, and that the ones at the N-terminus may function as targeting sequences in addition to other activities. Importantly, CTPs are highly polymorphic and accumulate in various locations, which suggests that they may have different activities. One way to proceed further in elucidating CTP1’s function will be to evaluate the relevance of plant protein interactors identified previously (Petre *et al.*, 2015).

## EXPERIMENTAL PROCEDURES

### Biological material and growth conditions

*Escherichia coli* (Subcloning Efficiency DH5*α* Competent Cells; Invitrogen, Carlsbad, California, USA) and *Agrobacterium tumefaciens* (electrocompetent strain GV3101) were conserved at -80°C and grown in L medium at 37°C and 28°C, respectively. *N. benthamiana* plants were grown in greenhouses at 22°C under 16/8h day/night conditions. The poplar hybrid Beaupré (*P. trichocarpa* × *Populus deltoides*) was propagated in greenhouse as previously described (Rinaldi *et al.*, 2007). Urediniospores of *Melampsora larici-populina* (isolate 98AG31) were conserved at -80°C and propagated as previously described (Rinaldi *et al.*, 2007). Beaupré leaf infections by *M. larici-populina* (isolate 98AG31, virulent on Beaupré) as well as *M. larici-populina* urediniospores germination *in vitro* were performed as previously described (Rinaldi *et al.*, 2007).

### Sequence analysis

Protein sequences were retrieved as follow: *M. larici-populina* (http://genomeportal.jgi-psf.org.html; Duplessis *et al.*, 2011), *M. lini* (Nemri *et al*, 2014), *Puccinia striiformis* f sp *tritici* and *Puccinia graminis* f sp *tritici* (Cantu *et al*, 2013), *N. benthamiana* (http://solgenomics.net/; *Bombarely et al.*, 2012) *and P. trichocarpa* (http://www.phytozome.net/poplar; *Tuskan et al.*, 2006). Protein sequence analyses were performed with ClustalX and Jalview softwares. Protein targeting sequences were predicted with SignalP 3.0 (http://www.cbs.dtu.dk/services/SignalP-3.0/), TargetP 1.1 (http://www.cbs.dtu.dk/services/TargetP/), ChloroP 1.1 (http://www.cbs.dtu.dk/services/ChloroP/) and Wolf P Sort (http://psort.hgc.jp/). Protein parameters were calculated with Protparam (http://web.expasy.org/protparam/).

### Cloning procedures and plasmids

The open reading frame (ORF) encoding CTP1_28-171_ and CTP2_29-185_ were amplified by polymerase chain reaction (PCR) using cDNA from Beaupré leaves infected by *M. larici-populina* (isolate 98AG31; Petre *et al.*, 2012), with primers designed to generate BbsI AATG/ggTTCG compatible overhangs. The ORF of CTP3_28-260_ was obtained through gene synthesis (Genewiz, London, UK), with codon optimization for plant expression and removal of internal BbsI and BsaI restriction sites. Sequences suitable for N-terminal tagging were generated by PCR amplifications with primers designed to insert BbsI AATG/GGCT compatible overhangs. Truncated sequences of CTP1 suitable for C-terminal tagging were generated by PCR amplifications with primers designed to insert BbsI AATG/ggTTCG compatible overhangs. DNA fragments were inserted into golden gate level 0 vectors pICSL01005 (BbsI compatible overhangs AATG/ggTTCG) or pICH41308 (BbsI compatible overhangs AATG/GGCT) following a digestion/ligation procedure (Weber *et al.*, 2011; Engler *et al.*, 2013; http://synbio.tsl.ac.uk/). Sequences of DNA fragments obtained by PCR were verified by sequencing. To generate C-terminally tagged proteins, the ORF of the protein of interest (BsaI compatible overhangs AATG/ggTTCG) was assembled with the ORF of a mCherry or a GFP (BsaI compatible overhangs ggTTCG/GCTT) into the golden gate level 1 binary vector pICH86988 (BsaI compatible overhangs AATG/GCTT; 35S promoter/OCS terminator). To generate N-terminally tagged proteins, the ORF of the protein of interest (BsaI compatible overhangs AATG/GCTT) was assembled with a 35S promoter (BsaI compatible overhangs GGAG/CCAT), the ORF of a mCherry or a GFP (BsaI compatible overhangs CCAT/AATG) and an octopine synthase (OCS) terminator (BsaI compatible overhangs GCTT/CGCT) into the golden gate level 1 vector pICH47742 (BsaI compatible overhangs GGAG/CGCT). Fusion protein sequences are shown in Table S1. Plasmids were multiplied using *E. coli*. Level 1 vectors were inserted in *A. tumefaciens* strain GV3101, and transformed bacteria were conserved at -80°C in 20% glycerol.

### Expression of proteins in *N. benthamiana*

A. *tumefaciens* strain GV3101 was used to deliver T-DNA constructs into leaf cells of three-week-old *N. benthamiana* plants, following the agroinfiltration method previously described (Win *et al.*, 2011). Briefly, overnight-grown bacterial cultures were adjusted at an OD_600_ of 0.2 into an infiltration buffer (10 mM MgCl_2_, 150 μM acetosyringone) one hour before leaf infiltration. For all co-transformations, *A. tumefaciens* strains were mixed in a 1:1 ratio in infiltration buffer to a final OD_600_ of 0.2. The leaves were collected two days after infiltration for further cell fractionation, protein isolation or microscopy. Stable *N. benthamiana* transgenics were obtained by A. tumefaciens mediated transformation (same strain as used for agroinfiltration), selected on Kanamycin and genotyped by PCR.

### Chloroplasts isolation

Three agrotransformed *N. benthamiana* leaves were collected two days after infiltration, cut into 1 cm^2^ pieces and incubated in 50 ml cold Isolation Buffer (IB; 400 mM Sorbitol, 50 mM HEPES KOH 8M, 2 mM EDTA) for 15 min. Leaf pieces were homogenized with a PT 1200 C Polytron homogenizer (3 × 5 sec full speed), and the lysate was immediately filtrated onto doubled Miracloth to remove cell debris. The filtrate was collected in 50 mL conical tubes and centrifuged at 3,000 rpm for 5 min at 4°C. The pellet containing the organelles was resuspended in 1 mL of cold IB and chloroplasts were isolated by centrifugation on a Percoll gradient (2 mL 80 % v/v Percoll/IB; 6 mL 40 % v/v Percoll/IB in a 15 mL conical tube) at 4,500 rpm for 10 min at 4°C. The lower green band corresponding to intact chloroplasts was collected by pipetting and immediately used for further experiments.

### Live-cell imaging by laser-scanning confocal microscopy

Small pieces of leaves (inferior face towards the objective) or solutions of isolated chloroplasts were mounted in water between a slide and a coverslip and immediately observed. Live-cell imaging was performed with a Leica DM6000B/TCS SP5 laser-scanning confocal microscope (Leica microsystems, Bucks, UK), using 10x (air) and 63x (water immersion) objectives. The GFP and the chlorophyll were excited at 488nm, the YFP was excited at 514 nm, and the mCherry was excited at 561 nm. Specific emission signals corresponding to the GFP, the YFP, the mCherry and the chlorophyll were collected between 505-525 nm, 530-550 nm, 580-620 nm and 680-710 nm, respectively, except where otherwise stated. Scanning was performed in sequential mode when needed. Image analysis was performed with Fiji (http://fiji.sc/Fiji).

### Protein isolation and western blotting experiments

*N. Benthamiana* leaves were harvested two days after infiltration, frozen in liquid nitrogen and grinded into powder with mortar and pestle. Total protein extraction was performed by reducing and denaturing proteins form the leaf powder 10 minutes at 95°C in Laemmli buffer (1 M pH 6.8 Tris-HCL, 10 mM dithiothreitol [DTT], 2% SDS, 20% glycerol) in order to avoid *in vitro* non-specific degradation of the fusion proteins. Then, five to twenty microliters of isolated proteins were separated by 15% SDS-PAGE, and protein content was estimated by coomassie blue staining. Western blotting experiments were performed as previously described (Pégeot *et al.*, 2014; Bozkurt *et al.*, 2014). The following antibodies were used: rabbit anti-GFP (Invitrogen), goat anti-rabbit IRW800 (LI-COR Biosciences) and GFP (B2): sc-9996 (Santa-Cruz Biotechnology). Membrane revelation was carried out on an Odyssey infrared imager (LI-COR Biosciences) or on an ImageQuant LAS 4000 luminescent imager (GE Healthcare Life Sciences).

## ACKNOWLEDGMENTS

We thank Y. Dagdas (TSL, Norwich, UK) and T. Bozkurt (Imperial College, London, UK) for critical reading of an early version of that manuscript. We thank P. Dodds (CSIRO, Canberra, Australia) for discussions. B. Petre was supported by an INRA Contrat Jeune Scientifique (CJS), by the European Union, in the framework of the Marie-Curie FP7 COFUND People Programme, through the award of an AgreenSkills’ fellowship (under grant agreement n° 267196) and by the laboratory of excellence (Labex) ARBRE, through the award of a mobility grant (12RW53). D. Saunders was supported by Leverhulme early career fellowship and a fellowship in computational biology at TGAC, in partnership with the John Innes Centre, and strategically supported by BBSRC. C. Lorrain is supported by an INRA CJS and by the Labex ARBRE. S. Duplessis is supported by the French National Research Agency through the Labex ARBRE (ANR-12-LABXARBRE-01) and the Young Scientist Grant POPRUST (ANR-2010-JCJC-1709-01). Research at TSL is supported by the Gatsby Charitable Foundation and the BBSRC.

---

**Figure S1.**
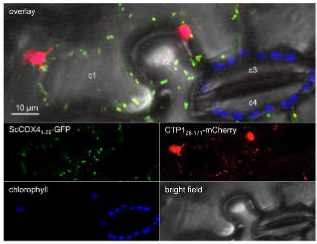
CTP1_28-171_-mCherry accumulates in mitochondria. Live-cell imaging of CTP1_28-171_-mCherry and ScCOX4_1-29_-GFP (mitochondria marker) in *N. benthamiana* leaf pavement cells. Proteins were expressed in *N. benthamiana* leaf cells by agroinfiltraion. Live-cell imaging was performed with a laser-scanning confocal microscope with a sequential scanning mode two days after infiltration. The GFP and the chlorophyll were excited at 488 nm; the mCherry was excited at 561 nm. GFP (green), mCherry (red) and chlorophyll (blue) fluorescent signals were collected at 505-525 nm, 580-620 nm and 680-700 nm, respectively. Images present a single optical section. Cells marked c1 and c2 are pavement cells differentially accumulating the two protein fusions. Cell marked c3 and c4 are guard cells.

**Figure S2.**
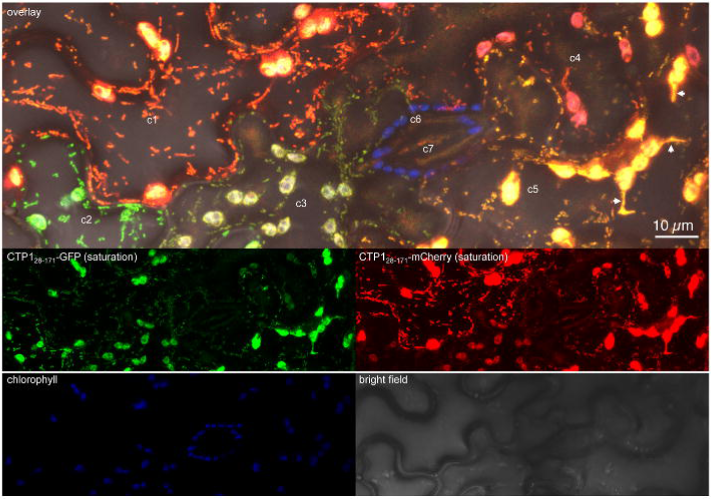
CTP1_28-171_-fluorescent protein fusions show no background in the cytosol. Same images as Figure 2, but with saturated GFP and mCherry fluorescent signals. See Figure 2 for details and annotations.

**Figure S3.**
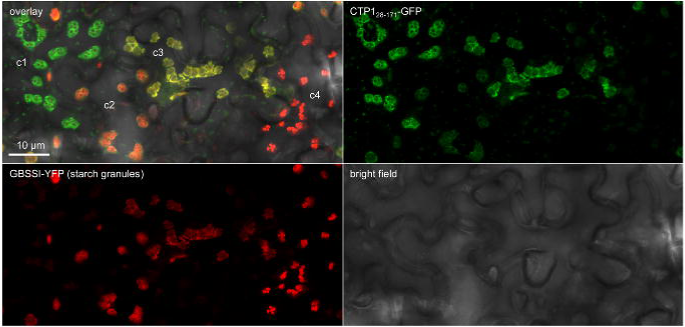
CTP1_28-171_-GFP diffuses in the stroma of chloroplasts. Live-cell imaging of CTP1_28-171_-GFP and GBSSI-YFP (starch granule marker) in *N. benthamiana* leaf pavement cells. Images present a maximal projection of 15 optical sections (z-stack: 12.0 μm). Proteins were expressed in *N. benthamiana* leaf cells by agroinfiltraion. Live-cell imaging was performed with a laser-scanning confocal microscope with a sequential scanning mode two days after infiltration. The GFP was excited at 488 nm; the YFP was excited at 514 nm. GFP (green) and YFP (yellow) fluorescent signals were collected at 505-510 nm and 530-550 nm, respectively. Cells marked c1 to c4 are pavement cells differentially accumulating the two fusion proteins.

**Figure S4.**
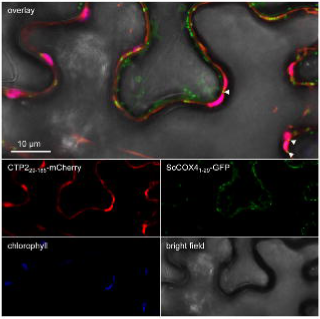
CTP2_29-185_-mCherry does not accumulate in mitochondria. Live-cell imaging of CTP2_29-185_-mCherry and ScCOX4_1-29_-GFP (mitochondria marker) in *N. benthamiana* leaf pavement cells. Images present a single optical section. White arrowheads indicate mitochondria in close vicinity to chloroplasts. Proteins were expressed in *N. benthamiana* leaf cells by agroinfiltration. Live-cell imaging was performed with a laser-scanning confocal microscope with a sequential scanning mode two days after infiltraion. The GFP and the chlorophyll were excited at 488 nm; the mCherry was excited at 561 nm. GFP (green), mCherry (red) and chlorophyll (blue) fluorescent signals were collected at 505-525 nm, 580-620 nm, and 680-700 nm, respectively.

**Figure S5.**
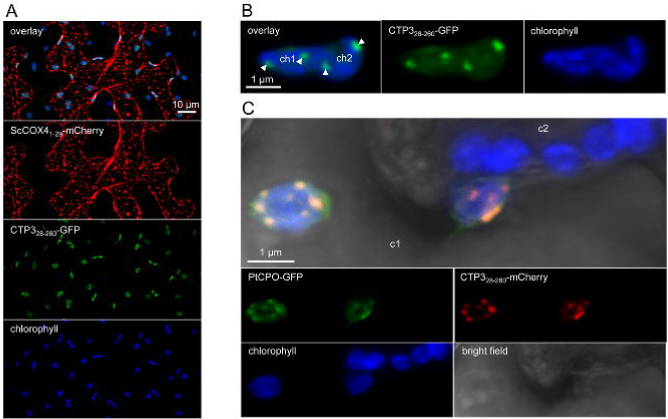
CTP3_28-260_-fluorescent protein fusions accumulate in discrete areas within chloroplasts and do not accumulate in mitochondria. (A) Live-cell imaging of CTP3_28-260_-GFP and ScCOX4_1-29_-mCherry (mitochondria marker) in *N. benthamiana* leaf pavement cells. Images present a maximal projection of 41 optical sections (z-stack: 32.8 μm). Due to the thickness of the Z-stack, the bright field is not interpretable and thus is not displayed). (B) Live-cell imaging of CTP3_28-260_-GFP in *N. benthamiana* leaf pavement cells. Images present a single optical section. ch1 and ch2 indicate two closely associated chloroplasts. White arrowheads indicate discrete areas inside chloroplasts that are labeled by CTP3_28-260_-GFP. (C) Live-cell imaging of CTP3_28-260_-mCherry and PtCPO (marker of undetermined discrete chloroplast bodies) in *N. benthamiana* leaf pavement cells. Images present a maximal projection of 11 optical sections (z-stack: 8.8 μm). The cell marked c1 accumulates both protein fusions, the cell marked c2 is a guard cell.

**Figure S6.**
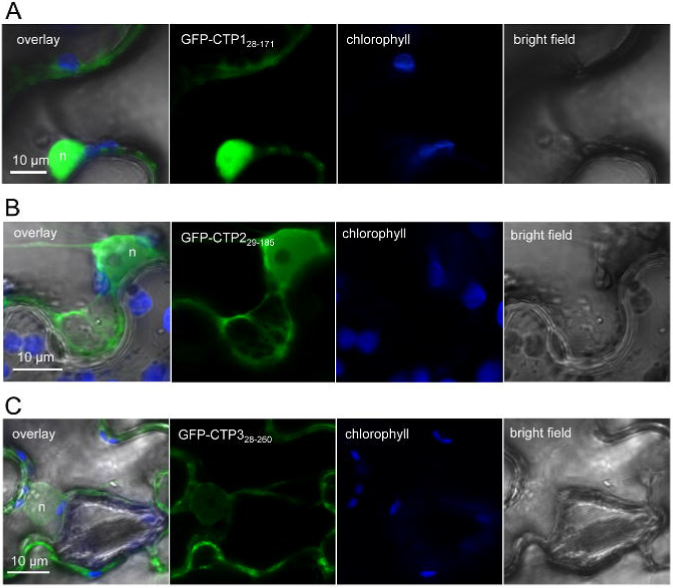
N-terminally tagged CTP1, CTP2 and CTP3 do not accumulate in chloroplasts. (A) Live-cell imaging of GFP-CTP1_28-171_ in *N. benthamiana* leaf pavement cells. Images show a single optical section. (B) Live-cell imaging of GFP-CTP2_29-185_ in *N. benthamiana* leaf pavement cells. Images show a maximal projection of three optical sections (z-stack: 2.4 μm). (C) Live-cell imaging of GFP-CTP3_28-260_ in *N. benthamiana* leaf pavement cells. Images show a single optical section. Live-cell imaging was performed with a laser-scanning confocal microscope two days after infiltration. The GFP and the chlorophyll were excited at 488 nm. GFP (green) and chlorophyll (blue) fluorescent signals were collected at 505-525 nm and 680-700 nm, respectively. n: nucleus.

